# How to achieve a higher selection plateau in forest tree breeding? Fostering Heterozygote x Homozygote relationships in Optimal Contribution Selection in the case study of Populus nigra

**DOI:** 10.1101/2021.02.15.431233

**Authors:** Mathieu Tiret, Marie Pégard, Leopoldo Sánchez

## Abstract

In breeding, Optimal Contribution Selection (OCS) is one of the most effective strategies to balance short- and long-term genetic responses, by maximizing genetic gain and minimizing global coancestry. Considering genetic diversity in the selection dynamic – through coancestry – is undoubtedly the reason for the success of OCS, as it avoids initial loss of favorable alleles. Originally formulated with the pedigree relationship matrix, global coancestry can nowadays be assessed with one of the possible formulations of the realized genomic relationship matrix. Most formulations were optimized for genomic evaluation, but few for the management of coancestry. We introduce here an alternative formulation specifically developed for Genomic OCS (GOCS), intended to better control heterozygous loci, and thus better account for Mendelian sampling. We simulated a multi-generation breeding program with mate allocation and under GOCS for twenty generations, solved with quadratic programming. With the case study of *Populus nigra*, we have shown that, although the dynamic was mainly determined by the trade-off between genetic gain and genetic diversity, better formulations of the genomic relationship matrix, especially those fostering individuals carrying multiple heterozygous loci, can lead to better short-term genetic gain and a higher selection plateau.

## 1 Introduction

Genomic selection (Meuwissen *et al*., 2001), in spite of using more precise Mendelian sampling terms compared to pedigree-based selection, and drastically increasing genetic response doing so, might accelerate the loss of genetic diversity and the useful variation per unit of time (Goddard, 2009; Hayes *et al*., 2009; Rutkoski *et al*., 2015). The loss of genetic diversity depends on a compromise between the co-selection of less relative individuals, which decreases the inbreeding rate per generation (Daetwyler *et al*., 2007), and the reduction of generation intervals, which gradually reduces the selective response (Grattapaglia, 2014). Yet, over the past decades, a growing number of authors have emphasized the importance of balancing loss of diversity and short-term genetic gain to sustain a long-term genetic response (Brisbane and Gibson, 1995; Woolliams *et al*., 2002; Jannink, 2010). This is especially true for perennial trees, for which minimal husbandry is provided while they can face harsh environments over a long period.

Optimal contribution selection (or OCS) was introduced by Meuwissen (1997) as a complement to the widely used Best Linear Unbiased Predictor (BLUP). BLUP conveniently integrates family information for increased accuracy, but also leads to rapid co-selection of relatives. On the other hand, OCS maximizes genetic gain while maintaining the inbreeding rate to a predefined level, thus accounting for the impact of genetic relationships on the population dynamics. OCS was originally formulated with the pedigree-based relationship matrix (A), but nowadays it can be easily adapted to the realized genomic relationship matrix (G). Its simplicity has made it today one of the most successful strategies to address the problem of selection-induced loss of diversity (Meuwissen *et al*., 2020). Furthermore, its flexibility has facilitated countless extensions. As a non-exhaustive illustration of its potential, De Beukelaer *et al*. (2017) extended OCS for different measures, such as heterozygosity, or the criterion of Li *et al*. (2008); Gebregiwergis *et al*. (2020) incorporated an alternative formulation of the genomic relationship matrix via QTL and markers; several studies proposed to combine OCS with mate selection, mainly to account for logistic constraints (*e.g*., the number of mating per individual), and to account for dominant effects (*e.g*., Varona and Misztal, 1999; Toro and Varona, 2010; Vitezica *et al*., 2013; Akdemir and Sánchez, 2016).

The realized genomic relationship matrix carries precise information on the relationship between pairs of individuals, but most of its formulations focused on shared homozygosity, which we could call homozygote x homozygote relationships. However, managing heterozygosity have a significant impact on the long-term genetic gain (De Beukelaer *et al*., 2017). Following the approach of Gebregiwergis *et al*. (2020), we propose here an alternative formulation of the genomic relationship matrix that focuses on individual carrying an excess of heterozygous loci relative to the population, or, in other words, the effect of individual-wise heterozygosity (for the study of population-wise heterozygosity, see De Beukelaer *et al*., 2017). Previous formulations of the realized genomic relationship matrix focused on increasing accuracy (e.g., Nejati-Javaremi *et al*., 1997; Fragomeni *et al*., 2017), or comparing genomic *vs* genealogical information (*e.g*., De Cara *et al*., 2011; Gomez-Romano *et al*., 2016). On the contrary, our method is intended to be used only in the context of Genomic OCS (GOCS), corresponding to the second genomic matrix in Gebregiwergis *et al*. (2020). Our study showed that developing different genomic relationship matrix for different usage can be beneficial, as our formulation increased long-term genetic gain when used in GOCS, but decreased accuracy when used in genomic evaluation.

In the present study, we will focus on heterozygote x homozygote relationships, and the objective is to exploit the impact of Mendelian sampling in the framework of genetic contribution (*e.g*., Avendaño *et al*., 2004; Cole *et al*., 2011). To do so, we devised a way to integrate heterozygote x homozygote relationship in the genomic relationship matrix, either to promote it or penalize it. We have developed a deterministic algorithm capable of solving both classical OCS and our alternative formulation, with quadratic programming. We have applied our method to the case study a population of *Populus nigra* L. (Salicaceae), to show by simulations that managing heterozygous loci in a multi-generation breeding program can achieve higher performances than with a classical OCS, and reach a higher Pareto curve. Among all the possible ways of constructing the genomic relationship matrix, the best strategy was to foster the heterozygote x homozygote relationship between pairs, which appeared to be the best compromise between genetic fixation and diversity.

## 2 Materials and Methods

### 2.1 Optimal contribution selection and convex optimization

Genetic contributions, first introduced by James and McBride (1958), are the cornerstone of OCS. The two opposing items in the optimization of OCS, gain and diversity, can be formulated according to the same decision variable, genetic contribution. The genetic contribution can be defined as the proportion of genes from a given ancestor that are present in a given cohort of descendants. More generally, the genetic contribution of an individual is its proportional contribution to the gene pool of the descendant population (Woolliams and Mäntysaari, 1995). In our case, when considering nonoverlapping generations, the genetic contribution of a given parent would simply be its proportional contribution to the offspring of the next generation. We denote *c* the vector of N genetic contributions, with N being the size of the parental population. Defined as such, ∑ *c*_i_ = 1.

With the knowledge of Y the vector of parental breeding values and A the numerator relationship matrix between parents (or G the realized genomic relationship matrix), and assuming no epistasis, it is possible to formulate the expected future performance or inbreeding coefficient of the population as *c*^T^Y (cY for simplicity) or as ½ *c*^T^A*c* respectively (or *c*^T^G*c* when using the genomic relationship matrix; cAc and cGc for simplicity). Deriving optimal selection decisions simultaneously accounting for future performance and future inbreeding is then possible through OCS, using genetic contributions as a decision variable. One of the possible formulations of such a problem is to solve min. λ cGc – cY, with λ a weighting parameter (see Woolliams *et al*., 2002 and references therein). It is important to note that the weighted average of cAc (or cGc) represents the expected inbreeding assuming *panmixia* or, more precisely, uniting in a full diallelic way all parents while respecting each parental *c*. This corresponds to the best expectation for inbreeding when the mating regime is unknown or not under the control of the breeder, which is often the case when, for example, mating is allowed to follow an open pollination regime as in a seed orchard.

As classical selection over cycles tends to accelerate the change of frequencies when they are intermediate, *i.e*., when the variances are at the highest levels, the risks of loss of favorable alleles by hitchhiking the alternative detrimentals would also increase (Sánchez *et al*., 2006). OCS would reduce this risk by maintaining frequencies at intermediate levels (when using an Identity-by-State matrix G, see Nejati-Javaremi *et al*., 1997), leading to a potentially slower fixation of favorable alleles, but overall benefits over the long-term genetic gain.

Originally, Meuwissen *et al*. (2001) formulated OCS as the maximization of genetic gain (cY) subject to a constrained inbreeding coefficient (cAc). Likewise, it is also possible to minimize the inbreeding rate while constraining the genetic gain (Akdemir and Sánchez, 2016). Choosing the adequate constraint is critical for populations never confronted to OCS. The methodology was primarily devised with long-term domesticated populations in mind, where records of change in inbreeding and genetic gain are typically known over several cycles, facilitating the choice of the constraints. In the absence of historical references, for novel species, a gradient of constraints would need to be evaluated *a priori*. In this sense, a holistic approach allowing visualization of the optimized function over a wide range of scenarios would be preferable as a start.

Following the approach of Akdemir *et al*. (2019), we considered OCS as a multi-objective optimization, where gain and diversity are improved simultaneously, i.e. maximizing gain and minimizing coancestry, by pondering weights that set the balance between the two items. The solutions of a multi-objective optimization, namely Pareto optima (Figure S1), delineate a twodimensional curve. Optimizing in two dimensions (genetic gain and coancestry) is mathematically equivalent to the scalarized version of the problem: minimize λ cGc - cY for any λ > 0. The scalarized problem has a unique global solution since the objective function is strictly convex (G is positive definite, as shown below). In other words, the curve of Pareto optima is the parametric curve, as a function of λ, of solutions minimizing λ cGc - cY. It is therefore the set of (c*Gc*, c*Y), where c* is the optimal contribution vector for a given λ. Without loss of generality, we consider the scalarized problem parameterized by α such that the problem becomes:

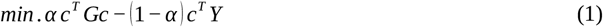

where α ∈ [0;1] can be interpreted as the trade-off value between coancestry and genetic gain (or the weight of coancestry compared to that of genetic gain).

Each OCS solution given a constraint (as in Meuwissen *et al*., 2001) is a particular Pareto optimum, or, in other words, is the solution of the scalarized problem for a particular α (if the constraint is not ill-formulated, *i.e*., not out of range). Both formulations – with α or with an inbreeding constraint – are strictly equivalent, and choosing a value for α is as arbitrary and difficult as choosing a value for a constraint without any *a priori*. Here, we will develop the framework with α, and different selection scenarios will be expressed in terms of α.

As in Meuwissen 1997, we have added some operational constraints to the multi-objective problem: the contributions must be larger than 0 to have a biological meaning but smaller than 0.5 to avoid selfing (0 ≤ c ≤ 0.5), and the sum of all contributions must be equal to 1 (**1**^T^c = 1). The constrained scalarized problem for a particular value of α is a constrained quadratic programming and can therefore be solved deterministically with an interior point method, adequate to solve constrained convex optimization (Boyd *et al*., 2004).

### 2.2 Different genomic matrices

Let X be the matrix describing the genotypes of the population, with L rows (number of markers) and N columns (number of individuals). The two homozygous states are encoded as −1 and 1, and the heterozygous state as 0 (as in VanRaden, 2008). We will consider the realized genomic relationship matrix formulated as G = X^T^X (N x N matrix). Note that the elements of X are not corrected by minor allele frequencies, nor is the resulting G scaled by the expected heterozygosity, as is usually done in genomic evaluation. Therefore, the diagonal elements of G provide information on the number of homozygous loci per individual, while the off-diagonal elements reflect the number of homozygous states shared by individuals across loci. Thus, for off-diagonals, the same homozygous state at a given locus adds one unit to the count, while one unit is subtracted for opposite homozygous states (not accounting for heterozygote by heterozygote, as pointed out by Gao and Martin, 2009), producing overall large values for pairs with resembling parents and small values for pairs with genetically distinct parents.

Using the G matrix defined above, OCS will penalize individuals with some particular relationship with the rest. For instance, an individual carrying multiple homozygous loci that are common in the population is less prone to be selected, as opposed to those carrying rare homozygous loci or heterozygous loci. In other words, per construction, individuals are scored depending on their homozygosity compared with other individuals’ homozygosity. We propose here to also score individuals on their heterozygosity compared with other individuals’ homozygosity; in other words, to account for the relative excess of heterozygous loci within an individual. The objective is to distinguish different Mendelian samplings occurring in identical homozygote x homozygote, which value in the G matrix is 1 (1 x 1 or −1 x −1 with the notation introduced above); opposite homozygote x homozygote, which value is −1 (−1 x 1); heterozygote x homozygote, which value is 0 (0 x 1 or 0 x - 1); and heterozygote x heterozygote, which value is also 0 (0 x 0). To distinguish heterozygote x homozygote (He x Ho), and heterozygote x heterozygote (He x He), we propose to change the value of He x Ho to β, which ranges from −1 to 1. A positive value of β means that few offspring will be generated from individuals with a large number of loci in He x Ho relationship with others. Hereafter, we denote G* the matrix constructed with β, where G* = G + β Q, Q being the matrix accounting for He x Ho loci, as G already account for homozygote loci (details in Supplementary materials).

In addition, we can derive that (details in Supplementary materials):

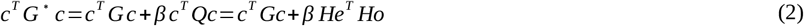

where He is the vector (of size L) of the proportion of heterozygous individuals contributing to the next generation for each locus, and Ho the equivalent for homozygous individuals. We can see from equation (2) that controlling β allows us to change the frequency of the carriers of favorable genotypic states, which in turn would favor the occurrence of certain crosses increasing the segregation of diversity, for instance by increasing the chance of a double heterozygous pairs over homozygous x heterozygous pairs. Such extra segregation could intuitively allow for a more sustainable genetic progress over the long term, without the risk of genetic hitchhiking.

For the objective function to be strictly convex, G* must be positive definite. Therefore, we performed a spectral projection of G* on the set of positive definite matrices, ensuring that the projected matrix is the closest positive definite matrix to G* (according to the spectral norm; Boyd *et al*., 2004). We will from here onward denote the projection of G* as G* itself to ease the reading.

### 2.3 Genomic data

The population used in the study included 1009 individuals from the French breeding population of *Populus nigra* (Pégard *et al*., 2020). All of them were genotyped with a 12k Infinium array (Faivre-Rampant *et al*., 2016) resulting in 5253 usable SNP markers after quality and frequency filtering (minor allele frequency higher than 0.05). The resulting genotypes were phased, imputed and a consensus recombination map derived (Pégard *et al*., 2019) by using FImpute software (Sargolzaei *et al*., 2014). The allelic effects were estimated from a genomic multitrait evaluation using breedR (Muñoz and Sánchez, 2020). In this study, we considered the trunk circumference as a focal trait, with a heritability of h^2^ = 0.5134 (Pégard *et al*., 2020). All the individuals were phenotyped.

Poplar is a dioecious species. Sex, however, cannot be determined before seven years of age, nor can it be predicted from the genomic profile yet. For this study, and to overcome the missing sex of unsexed candidates, we assumed a monoecious population. However, our method could be easily extended to dioecious populations by adding one simple constraint: d^T^c = 0.5, where d is a design vector indicating the female/male individuals.

The population *Populus nigra* was genetically structured (Figure S2), with a high linkage disequilibrium. In order to check for possible effects of the linkage, in addition to the original dataset, we shuffled the alleles at each locus so that the allele frequencies remained unchanged, while linkage disequilibrium could be partly removed. Hereafter, we will refer to it as “shuffled dataset”, as opposed to the original dataset that we will refer to as “unshuffled dataset”.

### 2.4 Simulation pipeline

We considered different simulation scenarios, assuming different values of α and β. We simulated multi-generation breeding schemes of a constant population size N at each generation, and the parameters (α and β) remained constant across generations within a given scenario. No introgression of external genetic material was considered here. As mentioned above, we estimated the allelic effects from the genotype data with a real phenotype (trunk circumference). The resulting allelic effects were considered “true” and constant across generations, and used to obtain the True Breeding Value (TBV) of the newly simulated candidates. Phenotypes and genotypes of all simulated individuals were considered to be known at each generation.

When simulating multi-generation breeding programs, the question arises as to how genomic estimated breeding values (GEBVs) should be assessed at each new virtual generation. With a heritability of h^2^ = 0.5134, environmental deviation was simulated as a normally distributed perturbation of mean 0 and variance (1 – h2)σ_g_^2^, where σ_g_^2^ was the TBV variance. At each generation, as was done in Jannink (2010) and De Beukelaer *et al*. (2017), GEBV was assessed with a ridge regression model (Searle 2006), using the matrix G (not G*). Genomic evaluation included all previous generations, so the reference population incrementally increased over time.

OCS was then applied at each generation on the simulated GEBVs, and the resulting contributions were converted into a mating plan that fits the OCS solution. We can notice that once the genetic contributions are fixed, the mating plan will not change the average breeding value, nor the average coancestry in the selected candidates. The mating, however, can change the progeny homozygosity – the diagonal elements of the matrix G at generation t+1 (Pryce *et al*., 2012; Sonesson *et al*., 2012). Expected progeny homozygosity is equal to parents’ genomic relatedness (up to a constant). To optimize the complementation of parents in mating, we computed the mating plan by minimizing the average progeny homozygosity per mating, *i.e*., the trace of the matrix G at generation t+1, with a linear programming (see Supplementary materials). Mate allocation was done with the transformed matrix G*. In order to assess the effect of mate allocation, random mating was also simulated. Finally, with the mating plan, we simulated the next generation with an *ad hoc* program (written in C++17). All scenarios started with the same initial population: the initial average breeding value was equal to 27.7 ± 25.9 and the population coancestry was equal to 0.214 ± 0.086.

Every simulated replicate proceeded for 20 (non-overlapping) generations, and was devised without mutation. The simulation followed a grid of parameters: α ranged from 0.1 to 0.9 (with steps of 0.1), and β was equal to −0.5, 0, or 0.5. Each parametric setting was simulated 100 times. In order to compare the results from different β, we also computed for each simulation the “true” coancestry, which is the coancestry the population would have had if β was equal to 0 (in line with the methods of Gebregiwergis *et al*., 2020). The whole pipeline was coded in R – 3.6.3 (R Core Team, 2020), and is available on github (https://github.com/mtiret/ocs.git).

### 2.5 Statistical analyses

Finally, in order to determine the relative importance of the factors α, β and mate allocation on genetic gain and true coancestry (through an analysis of variance) at a given generation, we considered the following statistical model:

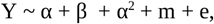

where Y is the output variable (either genetic gain or true coancestry), α and β the simulation parameters treated as fixed effects, α^2^ as the fixed effect of the squared parameter of α, m as the fixed effect of mate allocation (equal to 0 or 1), and e the residual fitted to a normal distribution. We considered here a polynomial regression (quadratic term α^2^) as there were symmetrical boundary effects of α upon Y: adding the quadratic term resulted in a gain of up to 70% in terms of adjusted coefficient of determination (r^2^). Quadratic α increases with a slower rate than linear α, hence the quadratic term can be interpreted as an effect that is constant for low values of α, but highly variable among high values of α. We will hereafter refer to this model as model (1). In some cases however, we focused on a given value of α, where we considered the following model:

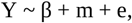

with equal notations and assumptions as model (1). We will hereafter refer to this model as model (2). These two models were analyzed using R - 3.6.3 (functions *lm* and *anova* from the base R package *stats*). We performed a type I analysis of variance, and the statistical significance of each factor was assessed with a Fisher’s *F*-test.

## 3 Results

### 3.1 Overall evolution of genetic gain and coancestry met common expectations

When considering the unshuffled dataset, only studying the effect of α (representing the weight of coancestry compared to that of genetic gain), and considering β (representing the value of He x Ho relationships) equal to 0 – corresponding then to the classical formulation of OCS – as expected, the average value of genetic gain increased for the whole period of selection, especially for low α (Figure 1, Table 1). True coancestry increased over time, but less so with higher α – as expected, when focusing on coancestry, genetic diversity increased over time. For the largest value of α (0.9), a consistent decrease was obtained for the whole period of selection. In other words, when α is high enough, though genetic gain is lower, producing genetic gain has no perceptible cost in terms of coancestry. De Cara *et al*. (2011) has already shown similar results, but is only possible when using genomic information (as opposed to pedigree information).

**Figure 1.**
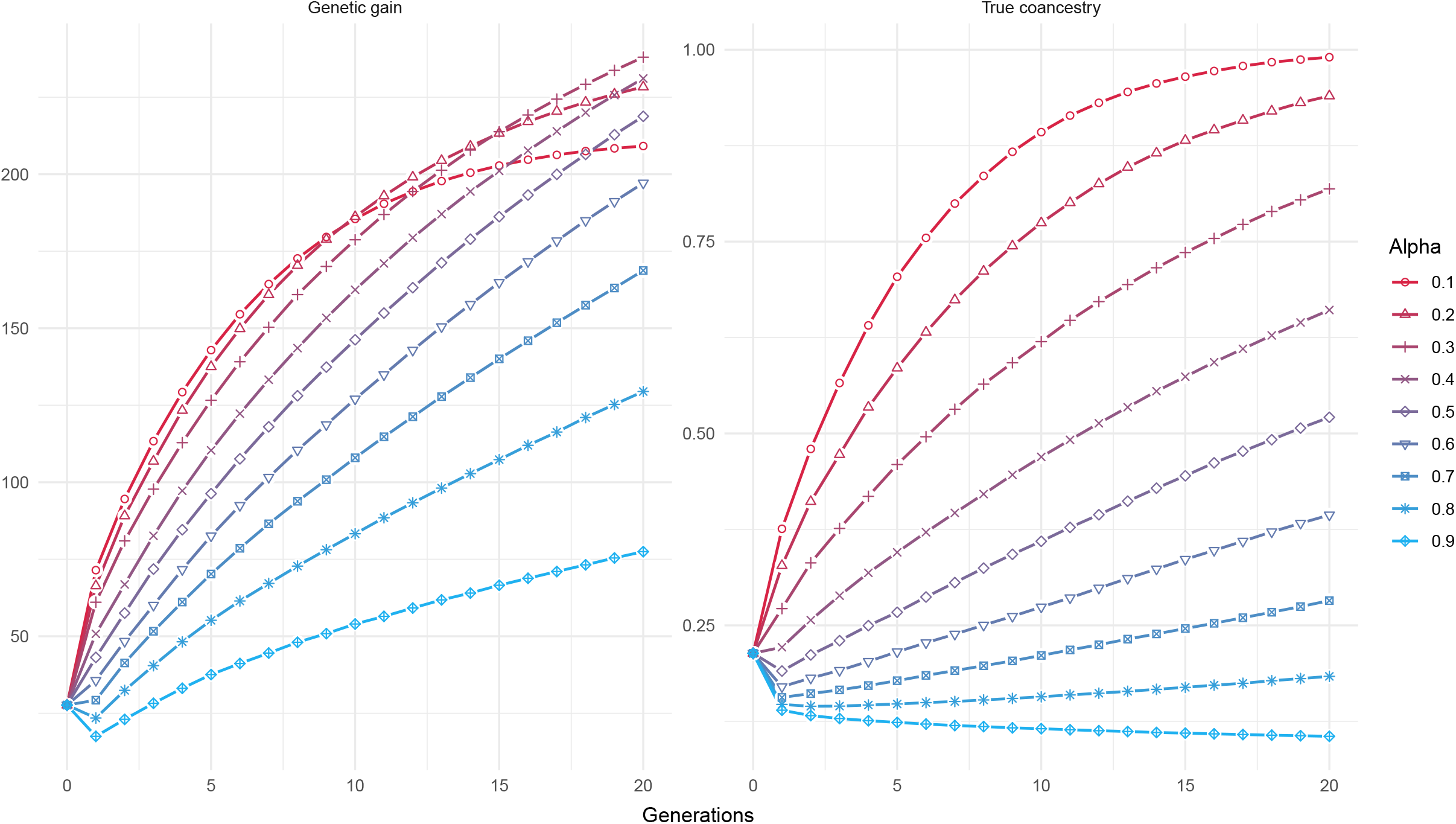
Evolution over time of genetic gain (left panel) and true coancestry (right panel), for different values of α (red for α = 0.1 till blue for α = 0.9), with β = 0, with the unshuffled dataset and h^2^ = 0.5134 (trunk circumference).

**Table 1.** Average values and standard deviations of genetic gain or true coancestry for trunk circumference (h^2^ = 0.5134) at generation 20, for the unshuffled dataset (S=0) and shuffled dataset (S=1), and without mate allocation (M=0) or with mate allocation (M=1). The initial states of genetic gain was 27.7 ± 25.9, and coancestry was 0.214.

|  |  | Genetic gain |  |  |  | True Coancestry |  |  |  |
| --- | --- | --- | --- | --- | --- | --- | --- | --- | --- |
|  |  | S = 0 |  | S = 1 |  | S=0 |  | S= 1 |  |
|  |  | M = 0 | M = 1 | M = 0 | M = 1 | M = 0 | M = 1 | M = 0 | M = 1 |
| $\alpha = 0.1$ | $\beta = -0.5$ | 208.1 ±<br>12.3 | 210.88<br>± 11.7 | 123.2<br>± 6.6 | 124.3<br>± 5.6 | 0.988 ±<br>0.005 | 0.988 ±<br>0.007 | 0.985 ±<br>0.006 | 0.984 ±<br>0.007 |
| | $\beta = 0$ | 207.1 ±<br>13.3 | 208.4<br>± 10.5 | 123.1<br>± 6.1 | 123.8<br>± 7.1 | 0.988 ±<br>0.006 | 0.990 ±<br>0.005 | 0.987 ±<br>0.006 | 0.986 ±<br>0.007 |
| | $\beta = 0.5$ | 204.7 ±<br>12.5 | 203.3<br>± 15 | 121.5<br>± 5.4 | 119.2<br>± 4.7 | 0.992 ±<br>0.004 | 0.992 ±<br>0.004 | 0.989 ±<br>0.006 | 0.990 ±<br>0.005 |
| $\alpha = 0.9$ | $\beta = -0.5$ | 146.4 ±<br>2.0 | 150.3<br>± 1.6 | 113.7<br>± 1.3 | 115.3<br>± 1.1 | 0.225 ±<br>0.001 | 0.227 ±<br>0.001 | 0.229 ±<br>0.001 | 0.230 ±<br>0.001 |
| | $\beta = 0$ | 74.5 ±<br>2.1 | 77.2 ±<br>2.1 | 93.3 ±<br>1.9 | 93.5 ±<br>2 | 0.107 ±<br>0.002 | 0.105 ±<br>0.002 | 0.181 ±<br>0.003 | 0.179 ±<br>0.003 |
| | $\beta = 0.5$ | 59.1 ±<br>2.7 | 60.8 ±<br>2.6 | 77.8 ±<br>2.7 | 77.3 ±<br>2.3 | 0.115 ±<br>0.003 | 0.103 ±<br>0.002 | 0.194 ±<br>0.004 | 0.173 ±<br>0.003 |

There is, however, no expectation regarding the genetic gain and coancestry variances, since OCS formulates its objective function and constraints in terms of expected values, not of variances. Compared to the initial standard deviation of genetic gain (25.9), simulations showed a consistent decrease over time (Table 1). Surprisingly, the decrease in standard deviation was even more pronounced for high α, but the coefficient of variation was lower. The same pattern was observed for true coancestry, where the standard deviation decreased in a more pronounced way for high α, but with a lower coefficient of variation. The lower coefficient of variation for both genetic gain and coancestry for high α suggests that the risk associated with a targeted genetic gain was better controlled when restricting coancestry, likely due to lower drift.

Different long-term horizons of genetic gain were reached depending on the value of α (Figure 1). In the relatively short-term, less than 6 generations, the achieved genetic gain decreased with increasing α. Eventually, in the longer term, the optimum α for genetic gain shifted from lower values to intermediate (at generation 20) and then to higher values of α (when reaching the selection plateau; Figure S3). The lower the value of α, the sooner the bend marking the start of the plateau. Such a plateau has already been described in the literature (Jannink, 2010; De Beukelaer *et al*., 2017), indicating a trade-off between the short and long-term horizons when setting the importance of gain versus diversity (α).

On the opposite, the optimum α for coancestry (i.e. α = 1) did not change across generations, *i.e*., the same ranking of α according to coancestries occurred for all generations. The most relevant feature, however, is that the change in coancestry between extreme values of α was larger than those observed for gain. In the long term (at generation 20), coancestry was multiplied by 10.23 when α shifted from 0.9 to 0.1, whereas for the same shift of α, genetic gain was only multiplied by 1.28. Such a difference in scale of response between gain and coancestry is clear even at the first generation of application of OCS. This trend can also be observed in the Pareto curve, where a substantial reduction in coancestry can have a minimum cost in gain whenever α is set between zero and intermediate values (α < 0.5; Figure S1).

### 3.2 The effect of α as the most explanatory factor after a long run

Statistical analyses of model (1) showed strong adjusted coefficients of determination for both genetic gain (unshuffled: r^2^ = 0.91; shuffled: r^2^ = 0.92) and coancestry (unshuffled: r^2^ = 0.98; shuffled: r^2^ = 0.99), suggesting a strong explanatory power of the parameters α, β and mate allocation. Overall, for both unshuffled and shuffled datasets, α was the most explanatory factor, with more than 84.6% of genetic gain variance explained, and more than 97.6% of coancestry variance explained (Table 2). When the dataset is not shuffled, genetic gain was mainly explained by the linear term of α, whereas when it is shuffled, genetic gain was mainly explained by the quadratic term. This result suggests that when linkage disequilibrium is not fully broken (unshuffled), every bit of increase in α has a perceptible effect on genetic gain, whereas for a broken linkage disequilibrium (shuffled), low α does not have any effect on genetic gain, only high values do. Coancestry, on the opposite, is always fully explained by the linear term, suggesting that although the effect of α is not always perceptible for genetic gain, it always is for coancestry. Mate allocation did only explain 0.0877% of the genetic gain variance in the unshuffled dataset, and was not significant in the shuffled dataset, suggesting that mating (defined as the minimization of progeny homozygosity) had only a small effect on genetic gain. Likewise, mate allocation only explained 0.00485% of the variance of coancestry in the unshuffled dataset, and only 0.00842% in the shuffled dataset.

**Table 2.** Statistical analysis (model (1)) of the simulations for trunk circumference (h^2^=0.5134), for the unshuffled (S = 0) and the shuffled (S = 1) datasets, for the genetic gain and true coancestry, at generation 20. Estimate are from model (1), and *p*-values from the analysis of variance. Significance levels are: *p*\*\*\* < 0.001; *p*\*\* < 0.01; *p*\* < 0.05; *p^n.s.^* > 0.05. ‘% Variance’: percentage of variance explained. r^2^: adjusted coefficient of determination of model (1).

| Effect | Genetic gain |  |  |  | True coancestry |  |  |  |
| --- | --- | --- | --- | --- | --- | --- | --- | --- |
| | S = 0 ($r^2 = 0.91$) | | S = 1 ($r^2 = 0.92$) | | S = 0 ($r^2 = 0.98$) | | S = 1 ($r^2 = 0.99$) | |
|  | Estimate | % Variance | Estimate | % Variance | Estimate | % Variance | Estimate | % Variance |
| $\alpha$ | 267*** | 58.7 | 276*** | 15.5 | -1.26*** | 97.5 | -1.25*** | 98.9 |
| $\alpha^2$ | -415*** | 25.9 | -310*** | 72.7 | 0.151*** | 0.100 | 0.236*** | 0.302 |
| $\beta$ | -34.0*** | 6.49 | -11.9*** | 3.98 | -0.0514*** | 0.434 | -0.0153*** | 0.0470 |
| **M** | 3.35*** | 0.0877 | 0.679<sup>n.s.</sup> | 0.0195 | -0.00443<sup>n.s.</sup> | 0.00485 | -0.00527*** | 0.00842 |

Favoring individuals with He x Ho relationship with other individuals (negative β) increased both genetic gain and coancestry (Table 1). β explained a slightly more important proportion of genetic gain variance for unshuffled dataset (6.49%) than for shuffled dataset (3.98%), suggesting that high linkage disequilibrium can benefit from a management of heterozygous loci. For coancestry, β only explained 0.434% (unshuffled) and 0.0470% (shuffled) of the variance. In the unshuffled dataset, when analyzing the effect of β on genetic gain with a fixed value of α (model (2)), β was not significant for α = 0.1 (Fisher’s *F*-test, *p* = 0.177), but significant for α = 0.9 (Fisher’s *F*-test, *p* < 0.001); the same pattern was observed in the shuffled dataset. The fact that β cannot have an impact for low α was expected by construction, but its significance for high α pinpoints that managing heterozygous loci can be beneficial in terms of genetic gain when there is sufficient diversity. On the opposite, β had a significant effect on coancestry (Fisher’s *F*-test, all *p* < 0.01), for low and high values of α, in the unshuffled and shuffled dataset. Overall, genetic gain and coancestry were mostly determined by α, even if the parameter was defined at the OCS level, and so, being blind to mate allocation. This result highlights the strikingly consistency of predictions with the genetic contribution framework. Similar results have been shown in previous studies (see Clark *et al*., 2013).

### 3.3 Favoring He x Ho relationship as the most sustainable strategy

The effect of β after twenty generations on genetic gain and true coancestry were different according to α (Figure 2). In the unshuffled dataset, for the lowest values of α (< 0.2, i.e. promoting the maximization of genetic gain over the minimization of coancestry), both genetic gain and coancestry were strongly influenced by α, but only negligibly by β. At this horizon, the population is detached from the Pareto curve. However, for larger α (> 0.2, i.e. promoting the minimization of coancestry over the maximization of gain), favoring He x Ho relationship (β < 0) resulted in best performances, i.e. a higher genetic gain for a given value of true coancestry. In other words, favoring He x Ho relationship allows the population to be on a higher Pareto curve (Figure 2). In addition, although mate allocation was almost negligible compared with α, it slightly shifted the Pareto curve upwards. On the opposite, in the shuffled dataset, β = 0 and β = −0.5 are on the same Pareto curve, suggesting again that the management of heterozygous loci would mainly be beneficial for population with a high linkage disequilibrium.

**Figure 2.**
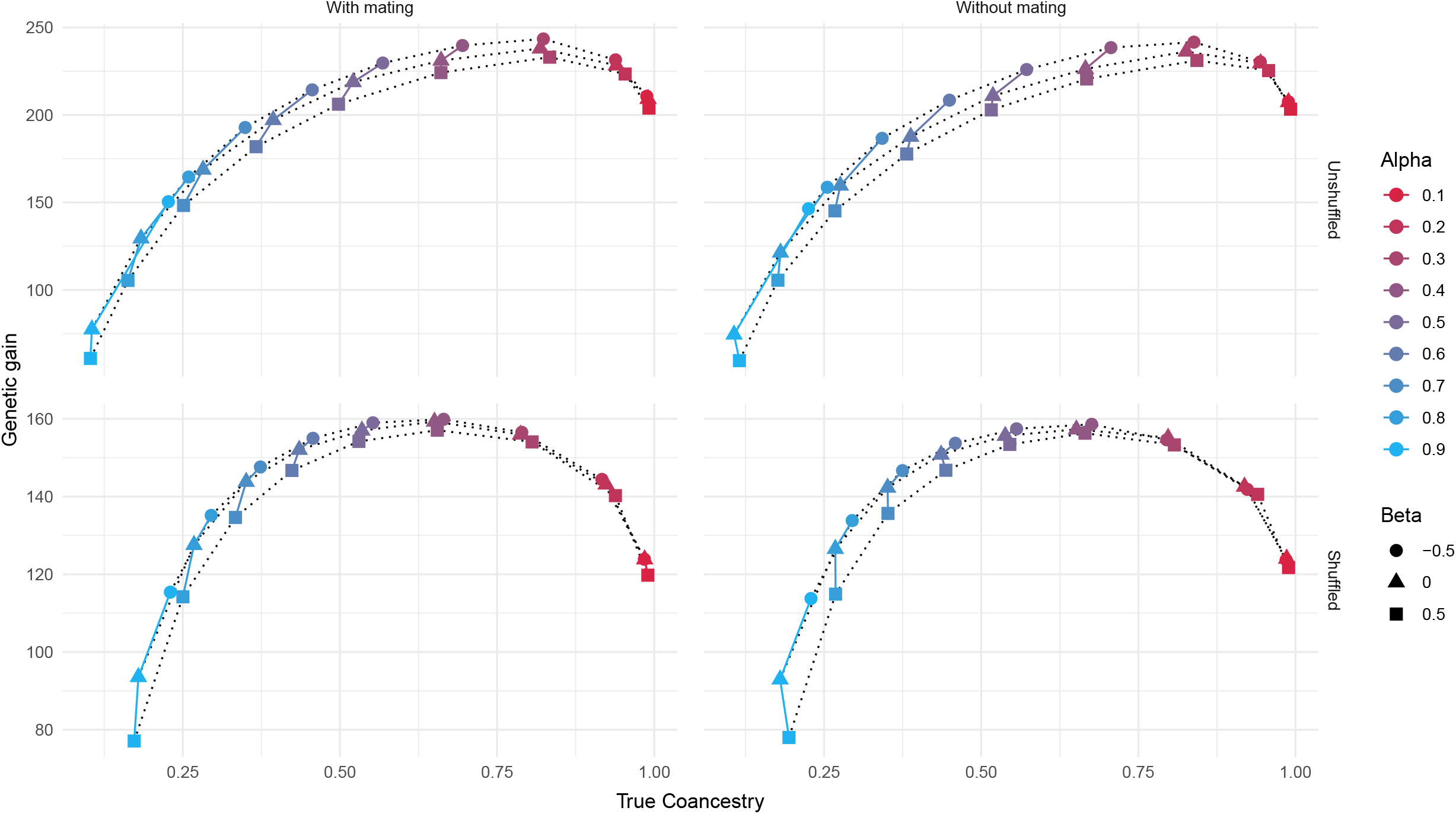
The Pareto optimum curve of true coancestry versus genetic gain, at generation 20, for different values of α (red for α = 0.1 till blue for α = 0.9) and β (circle for β = −0.5; triangle for β = 0; square for β = 0.5), with h^2^ = 0.5134 (trunk circumference). Left panels with mate allocation, right panels without mate allocation (random mating); top panels with the unshuffled dataset, bottom panels with the shuffled dataset.

Considering an alternative way of assessing the potential of a breeding program by measuring the GEBV of the population from favorable alleles that are not fixed yet, namely the breeding potential, the best strategy for the unshuffled dataset was to promote He x Ho relationship, especially for high values of α (Figure 3). Except for α = 0.1, the breeding potential of β = −0.5 was significantly higher than other values of β (Student’s *t*-test, all *p* < 0.05). For shuffled dataset however, the best strategy was to promote He x Ho relationship only for high values of α, as the breeding potential of β = −0.5 was significantly higher than other values of β only when α > 0.6 (Student’s *t*-test, all *p* < 0.05). The breeding potential of β = −0.5 and of a given α can almost reach that of a lower value of α. Therefore, favoring He x Ho relationship would increase short-term genetic gain, while guaranteeing a high selection plateau – as its level is determined by α, not β.

**Figure 3.**
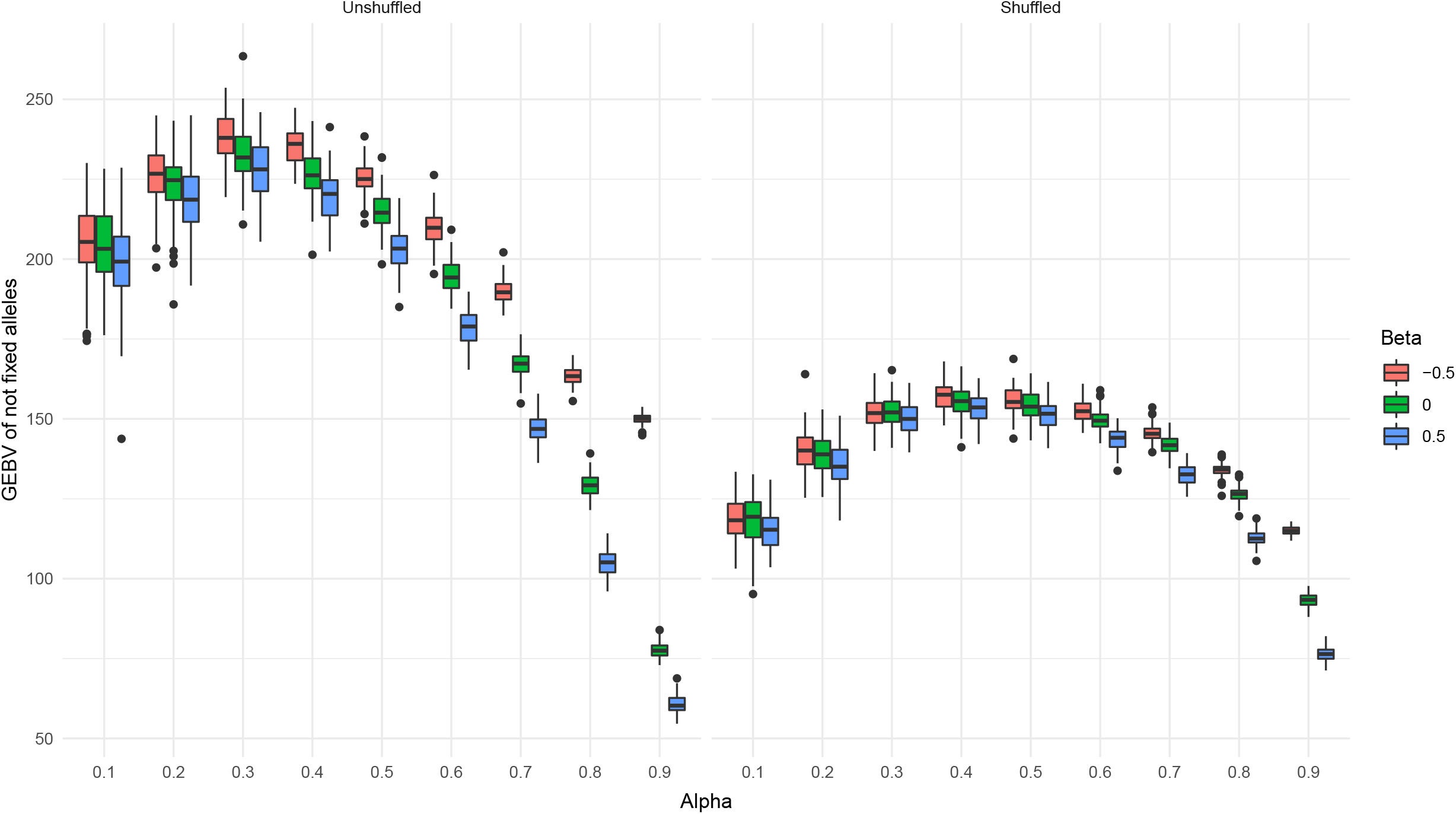
Boxplots of GEBV of not fixed alleles, at generation 20, for different values of α and β, with the unshuffled (right panel) and the shuffled (left panel) dataset, h2 = 0.5134 (trunk circumference).

## 4 Discussion

### 4.1 Long-term strategy in breeding programs

In a multi-generation breeding program, being able to select favorable alleles with little losses of favorable alleles on other loci would be the most desirable feature, *i.e*., avoiding the Bulmer effect (Bulmer, 1971). Drift often occurs through unwanted genetic hitchhiking when favorable and unfavorable alleles are trapped by limited sampling in continuous segments in linkage disequilibrium. There is therefore always a risk of loss as recombination might not be able to cope with the pace of selection and sampling generating the unfavorable linkage. One way to render recombination more efficient without slowing down the selection process would be to favor the pairing of candidates with a high potential for segregation in the offspring. It can be done more or less explicitly. One of the classical approaches, as shown in previous works (Jannink, 2010; De Beukelaer *et al*., 2017; Allier *et al*., 2019), consists in accounting for diversity through the trade-off parameter α, where diversity among candidates is modeled through relatedness or coancestry. This weighted approach, or the constrained formulation, can enhance the selection plateau, but often at the cost of slowing down the rate of progress. When such a perspective is applied to late maturing perennials, the cost in time for slow progress becomes a heavy drawback.

Another alternative to accelerate breeding without the drawback of drift is to minimize the uncertainties concerning the consequences of selection decisions, so that decisions can be based on sound predictions of the impact of selection on future generations. A way of doing this is to account for the He x Ho relationship between candidates, which is generally not considered in classical OCS. We have shown that optimizing mating, and modifying the genomic relationship matrix to account for He x Ho relationship, *i.e*., gain control over the Mendelian sampling, has always resulted in better performances, for both genetic gain and coancestry. The mating optimization was particularly important in the first generations, as already reported by Toro and Varona (2010).

When focusing on genetic gain (low α), better controlling Mendelian sampling had a small effect, meaning that the effect of drift cannot be counterbalanced. However, when considering higher α, controlling Mendelian sampling, especially by favoring He x Ho relationship (lower β) had a strong effect on the Pareto curve. Increasing genetic gain usually means: (i) a lower genetic (and genic) variance after selection, (ii) a higher level of fixation of favorable alleles, which constitutes the matter making up genetic gain, and (iii) a higher level of negative linkage disequilibrium covariance due to the Bulmer effect. The extra gain obtained from β < 0 could come from using more efficiently the genic variance, resulting in more depletion compared to that of higher β levels, and thus converting this available variation into favorable allele fixation, or likewise unfavorable allele elimination.

### 4.2 Alternative construction of genomic relationship matrix

Several studies proposed alternative formulations of the realized genomic relationship matrix (*e.g*., Nejati-Javaremi *et al*., 1997; VanRaden, 2008; Fragomeni *et al*., 2017) to improve accuracy in genomic evaluation, mainly assessing the efficiency of genomic information compared to genealogical information. Using alternative formulations might even lead to different long-term performance when used in the context of OCS (Gebregiwegis *et al*., 2020). However, a matrix that increases accuracy of genomic predictions does not necessarily measure inbreeding efficiently (Villanueva *et al*., 2021), so developing different formulations for different purposes seems preferable. Indeed genomic information has been used differently in different context (as reviewed in Maltecca *et al*., 2020): minimum coancestry mating (*e.g*., Fernández *et al*., 2021), selection against unfavorable alleles (*e.g*., Upperman *et al*., 2019), or genomic selection with dominance effects (*e.g*., Sun *et al*., 2014). In line with this recommendation, we developed a method specific to GOCS that showed consistently better long-term performances, although genomic evaluation with our alternative formulation (G*) showed a consistently lower accuracy than with G (5 to 10%, data not shown). In addition, when linkage disequilibrium was the highest (unshuffled dataset), our method performed best in terms of the Pareto curve.

Fostering the He x Ho relationships with the realized genomic relationship matrix might have the consequence of maintaining dominance effects, since it maintains potential for segregation. In addition to previous studies that showed the importance of accounting for dominance effects in genomic evaluation (*e.g*., Toro and Varona, 2010; Sun *et al*., 2013), our results show that a strategy that maintains dominance effects is also important for breeding programs. Consequently, we can argue that GOCS might benefit from including dominance effects in genomic evaluation, and optimize progeny homozygosity accordingly (Fernández *et al*., 2021). Whether dominance effects and its variance are only important to account for as a correction term in genomic evaluation (*e.g*., to improve accuracy), or have deeper implications in the population dynamic remains an open question.

### 4.3 Variance in OCS

The main challenge of OCS lies in the management of stochasticity: the objective function, as stated above, is formulated with expected values, and not with variances, thus neglecting the variability caused by the uncertainties of random mating and Mendelian sampling. This leads to stochasticity around the predicted Pareto optima, which analytical formula are only known for very few special cases (*e.g*., Garcia-Cortes *et al*., 2013). In our study, the undesired stochasticity was fixed by mate allocation, making segregation and environmental deviation the only sources of random variation. Mate allocation did, however, have a surprisingly low proportion of variance explained compared with what was reported previously (*e.g*., Hamrick and Godt, 1996; Nybom, 2004). The main challenge remains to efficiently convert the remaining additive variance (from segregation and environmental deviation) in future genetic gain (Santos *et al*., 2019), which can be done by integrating variance terms in the objective function.

Introducing variability in the parameter α and β could also be desirable in the case of multigeneration breeding, such as considering different values of α and β at each generation, depending on the current state of the population. For instance, considering a high value of α (high diversity) could be important at the very short-term to prevent losses of favorable alleles in low frequency, but once the Bulmer effect is absorbed by recombination, it could be safe to switch to a more aggressive strategy such as with lower values of α. Preliminary works showed indeed that a geometrically decreasing α along the generations with a rate of 0.99 brings better long-term performances. Differential selection over generation is therefore a field worth investigating, and warrants further studies.

### 4.4 Conclusion

In this article, we have extended OCS by proposing an alternative formulation of the realized genomic relationship matrix that better account for the Mendelian sampling. In multi-generation breeding programs it is important to account for diversity to reach a higher selection plateau, even though the speed at which it is reached can be slow. We have shown by and large that the population dynamic is overall dominated by the trade-off value α between genetic gain and genetic diversity. However, better accounting for Mendelian sampling, even implicitly as proposed here by fostering individuals with multiple loci in He x Ho relationship with others, could minimize the speed problem, by “accelerating” the breeding while maintaining a high level of diversity and selective potential for future generations.

## Acknowledgments

The authors acknowledge the GIS peuplier for access to data. This study was funded by the following sources: the INRA SELGEN funding program (project BreedToLast) providing sequencing, genotyping data and PhD grant for MP; and from the European Union’s Horizon 2020 research and innovation program under grant agreement No 676876 (GenTree) for basic functioning and postdoc grant for MT and MP.

## Data Availability statement

The data that support the findings of this study are openly available in DRYAD at https://doi.org/10.5061/dryad.0rxwdbs1k.

