## Supplementary material for "How to achieve a higher selection plateau in forest tree breeding? Fostering Heterozygote x Homozygote relationships in Optimal Contribution Selection in the case study of Populus nigra": supplementary.docx

Supplementary materials

#### Proof of equation (2)

Let X be the matrix describing the genotypes in the population, with L rows (number of markers) and N columns (number of individuals). The two homozygous states are encoded as −1 and 1, and the heterozygous state as 0 (as in Van Raden, 2008). Let then Q defined as follows:

Q = 0.5 L_NxN_ − 2 (|X| − ½_LxN_)^T^ (|X| − ½_LxN_),

where L_NxN_ a matrix of size N x N with each value equal to L, |X| being the matrix X with each value transformed to its absolute value, and ½_LxN_ a matrix of size L x N with each value equal to ½. This matrix Q is a matrix of size N x N, for which each value is the count of 0 x 1 or 0 x -1. Likewise, let W defined as follows:

W = ( (X − 1_LxN_) ၀ (X + 1_LxN_) )^T^ ( (X − 1_LxN_) ၀ (X + 1_LxN_) ),

with identical previous notation, and ၀ the Hadamard product. This matrix W is a matrix of size N x N, for which each value is the count of 0 x 0. Let now *c* be the genetic contribution, a vector of size N, for which each value is between 0 and 0.5 and for which the sum is equal to 1. Therefore, we have the following:

c^T^Qc = 0.5 L − 2 c^T^ (|X| − ½_LxN_)^T^ (|X| − ½_LxN_) c

= 0.5 L − 2 ∑_k_ ( ∑_i_ c_i_ (|x_k,i_| − ½) )^2^ = 0.5 L − 2 ∑_k_ (ho_k_ − ½)^2^, with hok the total contribution of homozygous individuals for the locus k,

= 0.5 L − 2 ∑_k_ (ho_k_^2^ − ho_k_ + ¼) = 2 ∑k ho_k_ (1 − ho_k_) = 2 Ho^T^ He,

where Ho is the vector of ho_k_, and He the equivalent for heterozygous individuals.

As for W, using similar reasoning and remarking that c ( (X − 1_LxN_) ၀ (X + 1_LxN_) ) is equal to He, we can deduce that :

c^T^Wc = He^T^ He.

Therefore, we can state that:

c^T^G*c = c^T^Gc + β Ho^T^He + γ He^T^He.

### Supplementary figures

**Figure S1.** The Pareto optimum graph : the x-axis is the coancestry of a population, and the y axis is the average genetic gain of a population. Each point is a simulation (one population), each color is a value of α (dark blue for α = 0.1, light blue for α = 0.9). The dotted line is the Pareto optimum prediction of OCS. Simulated with β = γ = 0, and h^2^ = 0.66.

**Figure S2.** Principal Component Analysis of the genotypes of the 1009 individuals of *Populus nigra*. Each point is an individual, and each color is the quartile of GEBVs (1 are the lowest GEBVs, 4 are the highest). The first two axes explained respectively 9.7% and 6.7% of the variance.

**Figure S3.** Estimate of the effect of β and γ in the model (2), for genetic gain, for different values of α, at generation 20 and for h^2^ = 1.

**Figure S4.** Proportion of sum of squares of the effects of β and γ in the model (2), for genetic gain, for different values of α, at generation 20 and for h^2^ = 1.

**Figure S5.** Estimate of the effect of β and γ in the model (2), for coancestry, for different values of α, at generation 20 and for h^2^ = 1.

**Figure S6.** Proportion of sum of squares of the effects of β and γ in the model (2), for coancestry, for different values of α, at generation 20 and for h^2^ = 1.

**Figure S7.** Genic variance for different values of β and α (dark blue for α = 0.1, light blue for α = 0.9), at generation 20 and for h^2^ = 1.

**Figure S8.** Example of genotypic covariance for different values of β (dark blue for β = -1, light blue for β = 1), with α = 0.8, γ = 0 and h^2^ = 1.

**Figure S9.** Selection efficiency for different values of β (dark blue for β = -1, light blue for β = 1), at generation 20 and with γ = 0 and h^2^ = 1. Selection efficiency is the ratio of the accumulated allelic effects due to selection (fixed favorable allelic effect + lost unfavorable allelic effect) and the accumulated allelic effects due to drift (fixed unfavorable allelic effect + lost favorable allelic effect).

### Supplementary tables

**Table S1.** Illustration of genomic relationship matrix for one locus, and with the -1, 0 and 1 encoding (as in VanRaden 2008).

| parents | − 1 | 0 | 1 |
| --- | --- | --- | --- |
| − 1 | 1 | 0 | − 1 |
| 0 | 0 | 0 | 0 |
| 1 | − 1 | 0 | 1 |

**Table S2.** Analysis of variance (type I) of the model (1), for genetic gain at generation 1 and 20, on the shuffled dataset with h^2^ = 1. Sum of squares is shown as a percentage of the total sum of squares. Significance levels are : *p**** < 0.001; *p** <* 0.01; *p** < 0.05; *p*^n.s.^ > 0.05. Adjusted coefficient of determination is denoted r^2^_adj_.

|  | t = 1 (r^2^_adj_ = 0.86) | | | t = 20 (r^2^_adj_ = 0.79) | | |
| --- | --- | --- | --- | --- | --- | --- |
|  | Sum of Squares (%) | Df | F value | Sum of Squares (%) | Df | F value |
| α | 76.8 | 1 | 125124*** | 21.0 | 1 | 22446*** |
| β | 4.0 | 1 | 6560*** | 15.9 | 1 | 16923*** |
| γ | 1.4 | 1 | 2325*** | 0.007 | 1 | 7.0** |
| α^2^ | 4.0 | 1 | 6448*** | 42.0 | 1 | 44769*** |

**Table S3.** Analysis of variance (type I) of the model (1), for coancestry at generation 1 and 20, on the shuffled dataset with h^2^ = 1. Sum of squares is shown as a percentage of the total sum of squares. Significance levels are : *p**** < 0.001; *p** <* 0.01; *p** < 0.05; *p*^n.s.^ > 0.05. Adjusted coefficient of determination is denoted r^2^_adj_.

|  | t = 1 (r^2^_adj_ = 0.95) | | | t = 20 (r^2^_adj_ = 0.93) | | |
| --- | --- | --- | --- | --- | --- | --- |
|  | Sum of Squares (%) | Df | F value | Sum of Squares (%) | Df | F value |
| α | 78.2 | 1 | 389011*** | 91.3 | 1 | 291339*** |
| β | 0.07 | 1 | 331*** | 1.4 | 1 | 4311*** |
| γ | 0.07 | 1 | 322*** | 0.01 | 1 | 317*** |
| α^2^ | 17.1 | 1 | 85233*** | 0.2 | 1 | 647*** |
