## Supplementary figures and images for "How to achieve a higher selection plateau in forest tree breeding? Fostering Heterozygote x Homozygote relationships in Optimal Contribution Selection in the case study of Populus nigra"

### figure_S1.png

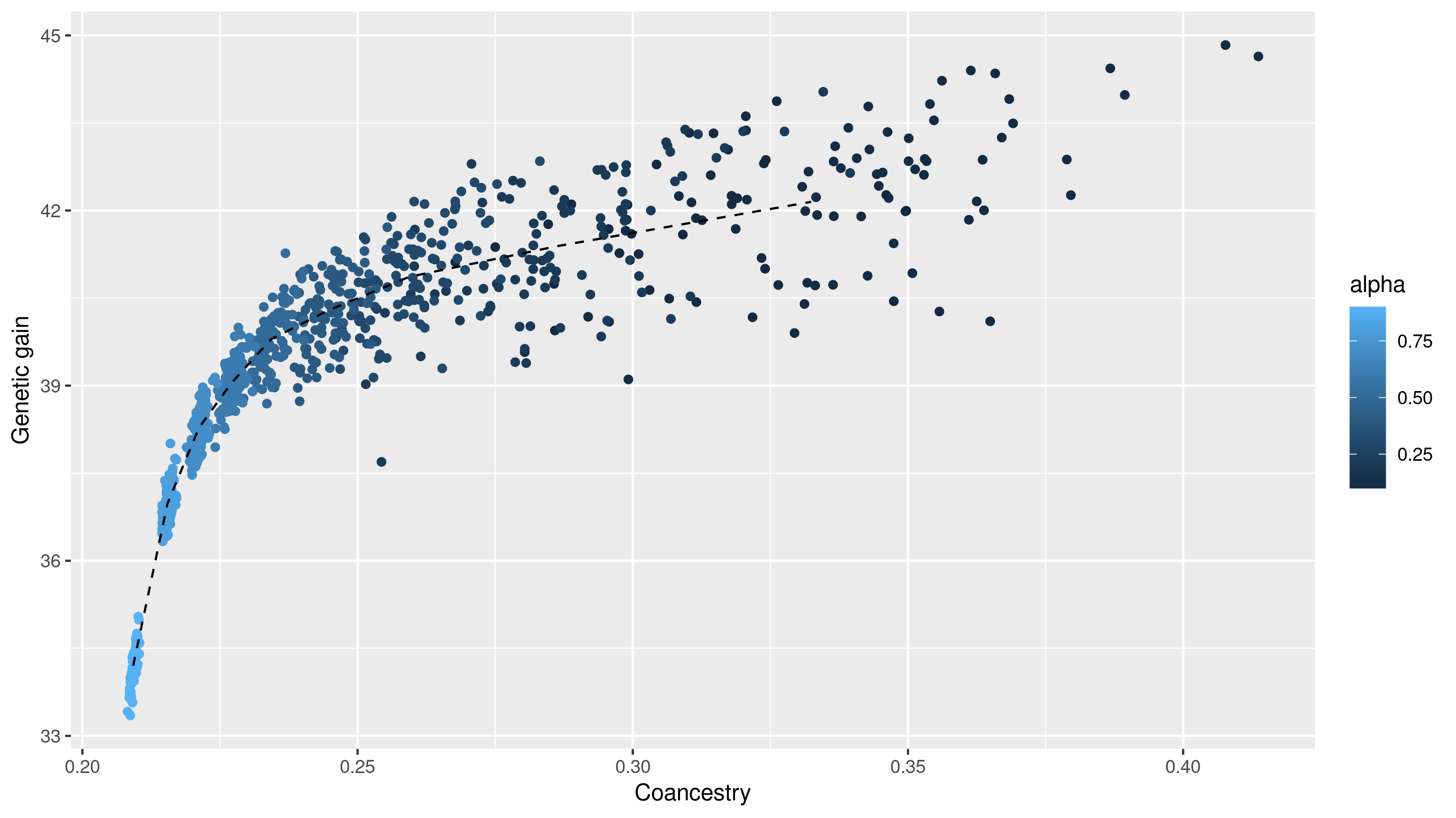

### figure_S2.png

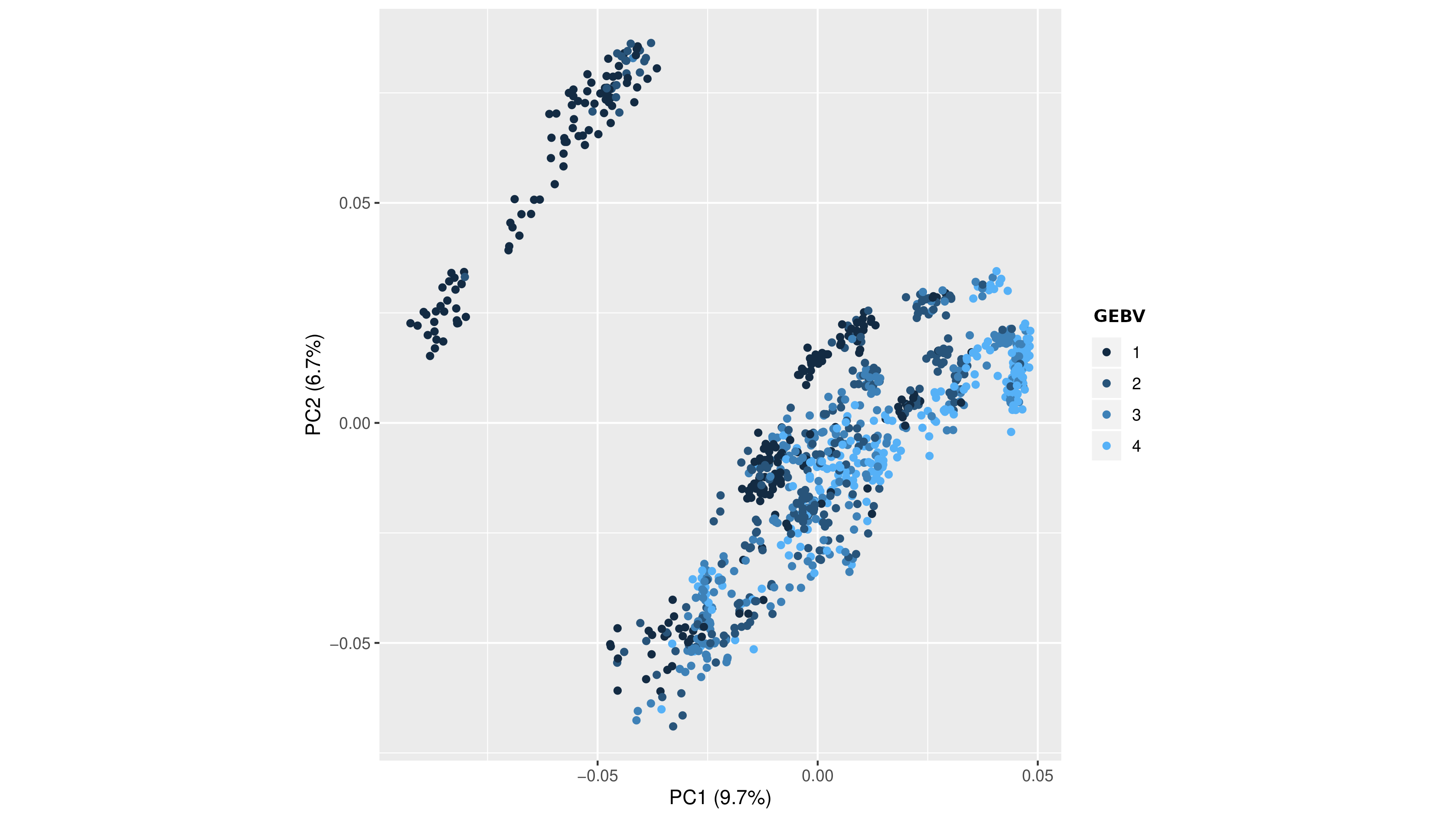

### figure_S3.png

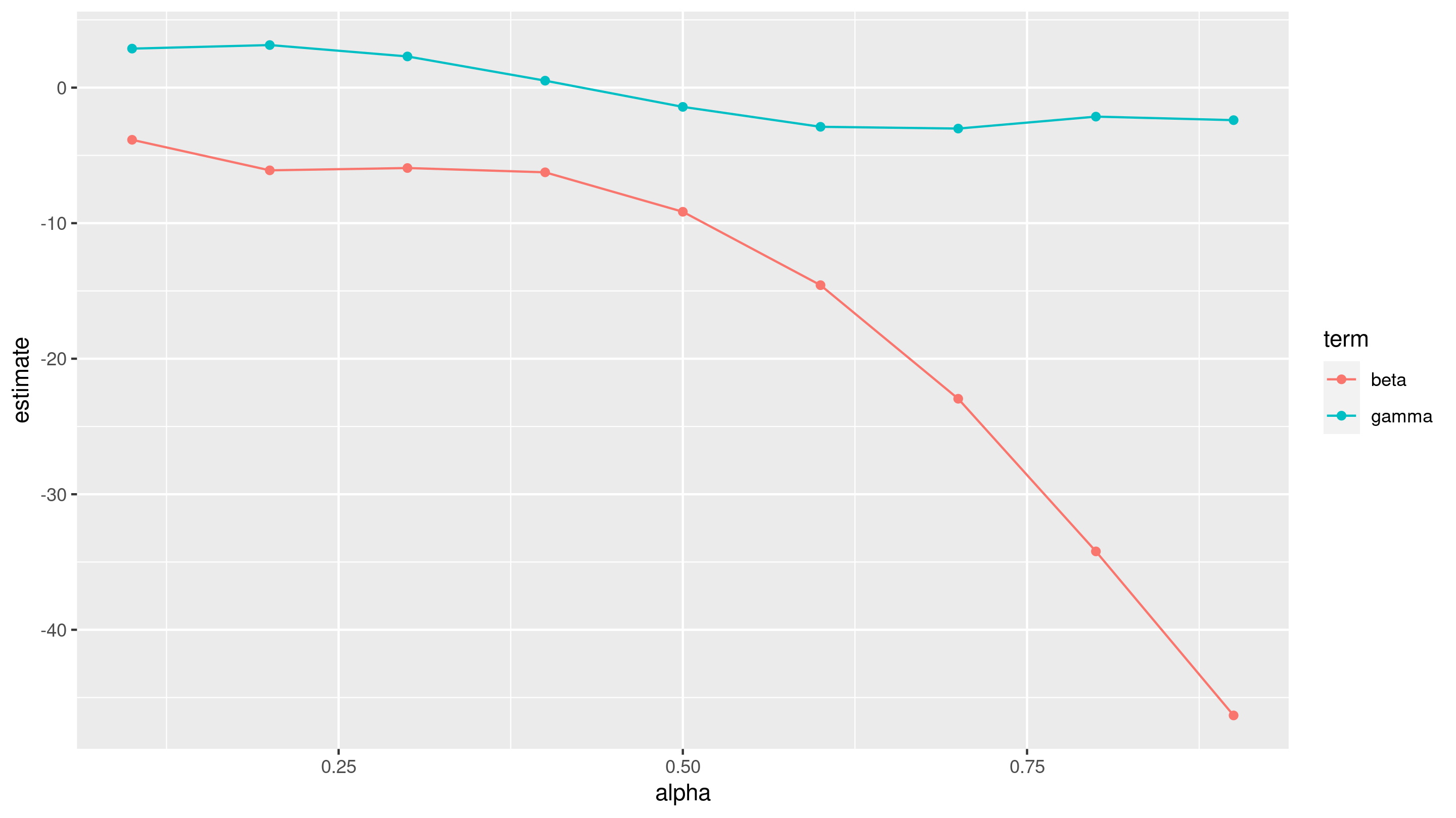

### figure_S4.png

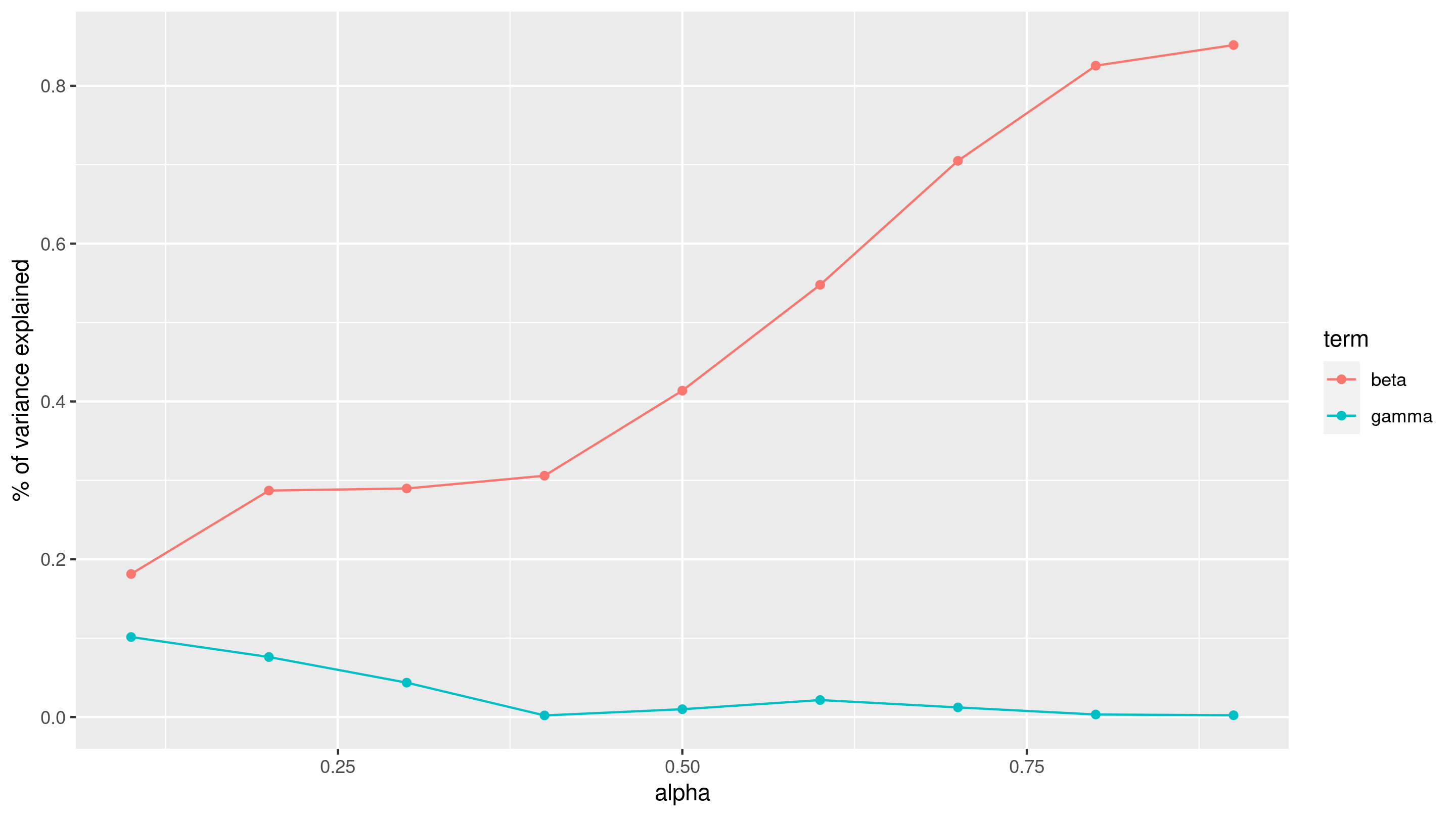

### figure_S5.png

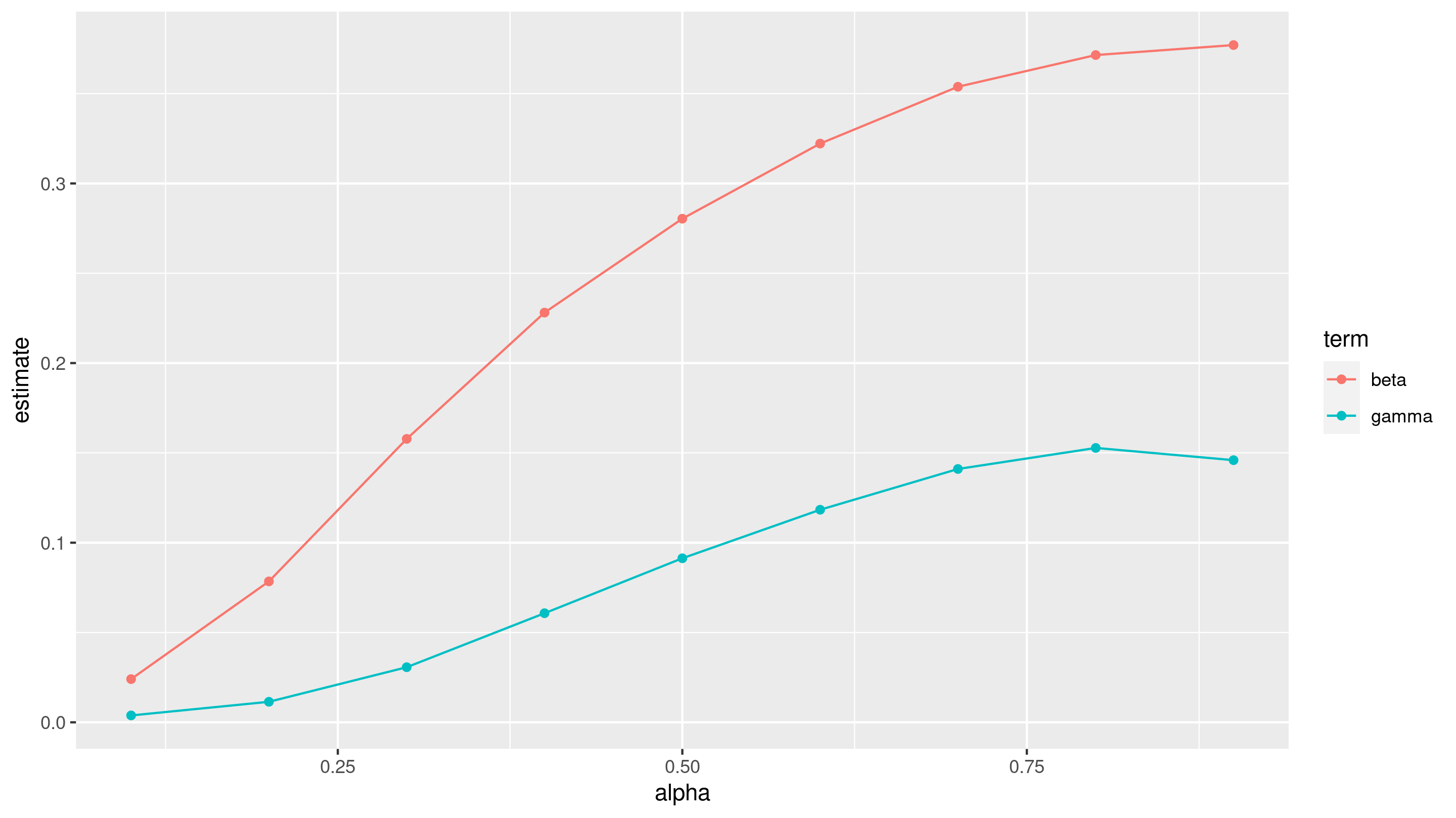

### figure_S6.png

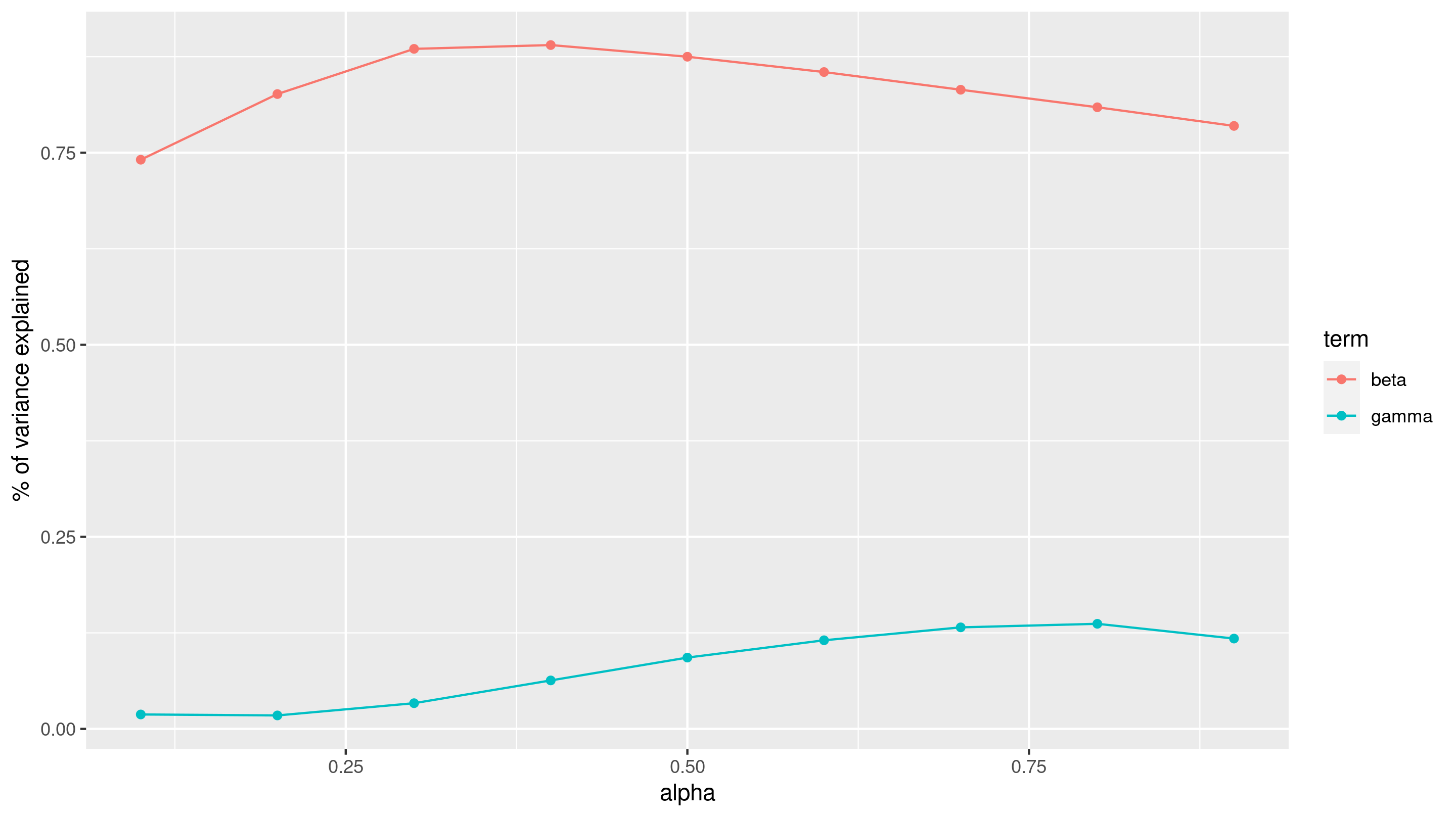

### figure_S7.png

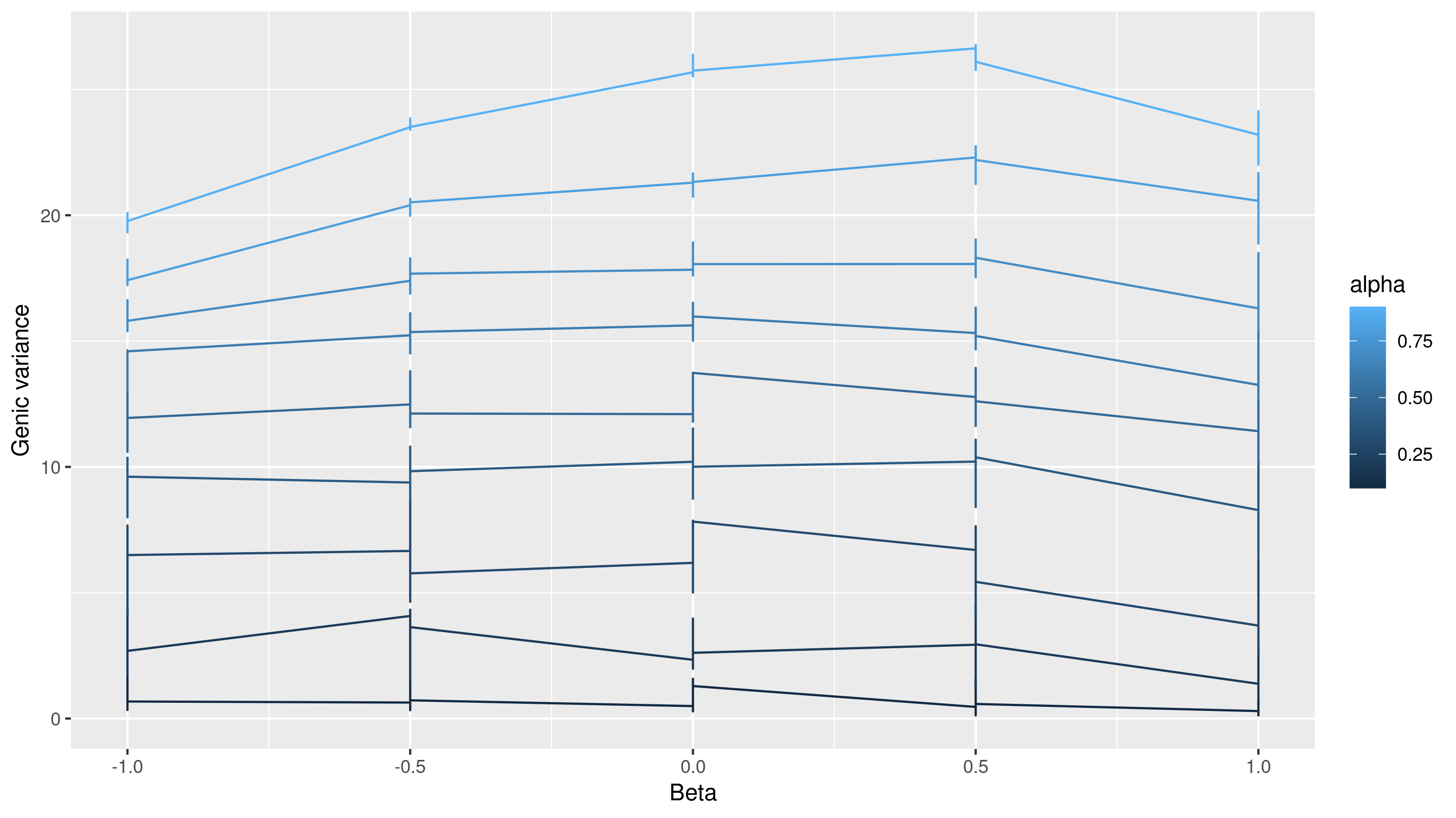

### figure_S8.png

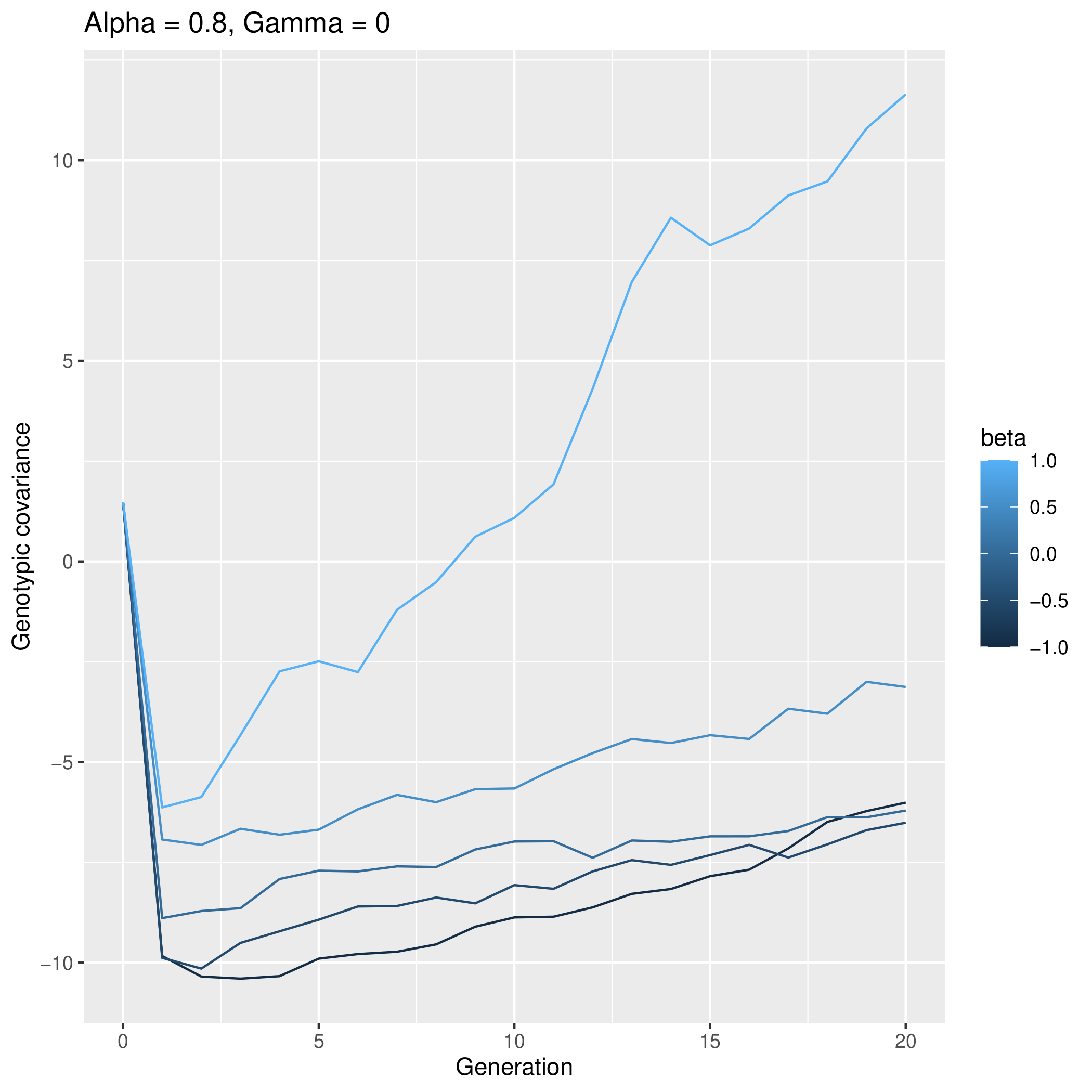

### figure_S9.png

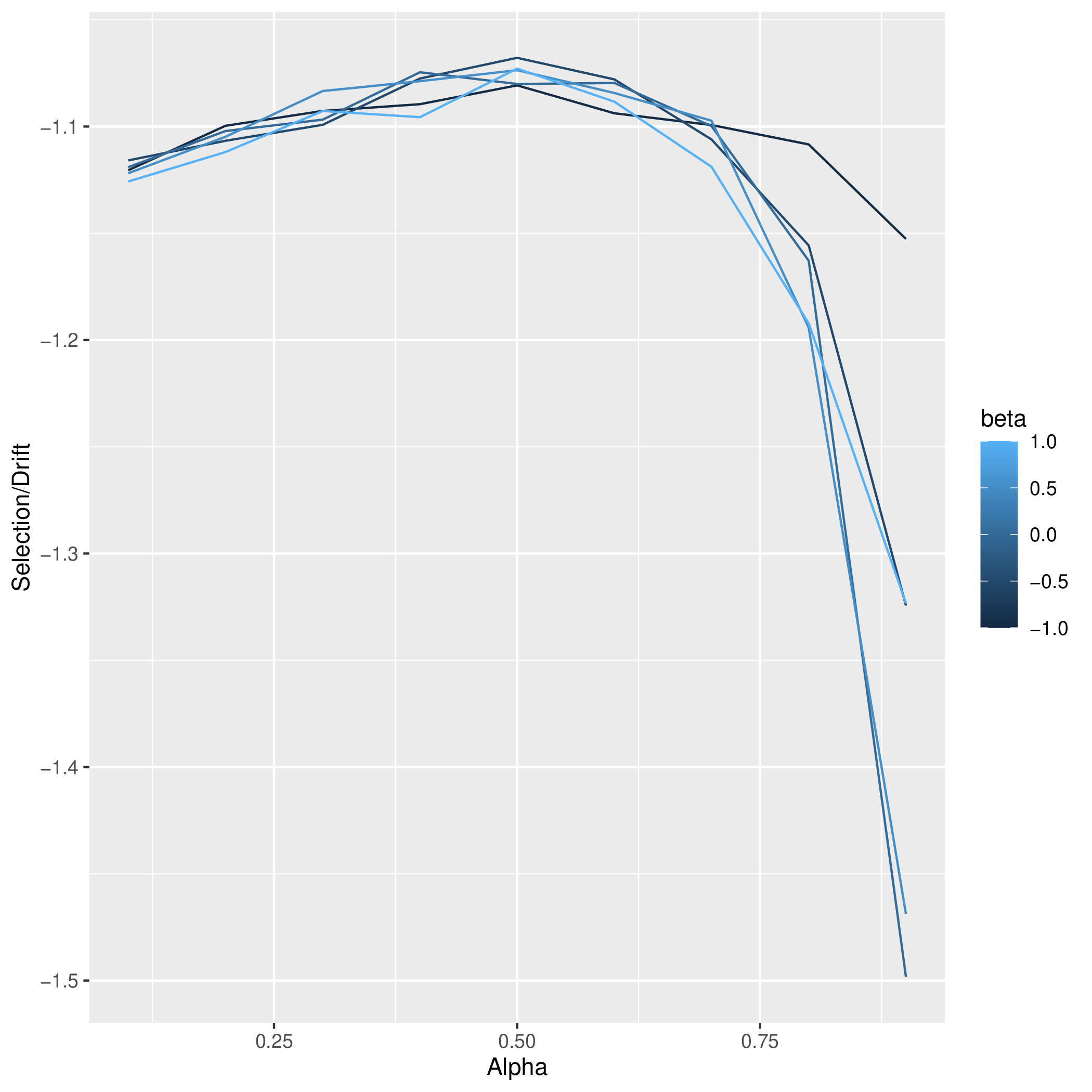
